# Genotype-phenotype association mining in bipolar disorder: market research meets complex genetics

**DOI:** 10.1101/116624

**Authors:** René Breuer, Manuel Mattheisen, Josef Frank, Bertram Krumm, Jens Treutlein, Layla Kassem, Jana Strohmaier, Stefan Herms, Thomas W. Mühleisen, Franziska Degenhardt, Sven Cichon, Markus M. Nöthen, George Karypis, Bipolar Disorder Genetics (BiGS) Consortium, Francis J. McMahon, Marcella Rietschel, Thomas G. Schulze

## Abstract

Disentangling the etiology of common, complex diseases is a major challenge in genetic research. For bipolar disorder (BD), several genome-wide association studies (GWAS) have been performed. Similar to other complex disorders, major breakthroughs in explaining the high heritability of BD through GWAS have remained elusive. To overcome this dilemma, genetic research into BD, has embraced a variety of strategies such as the formation of large consortia to increase sample size and sequencing approaches. Here we advocate a complementary approach making use of already existing GWAS data: applying a data mining procedure to identify yet undetected genotype-phenotype relationships. We adapted association rule mining, a data mining technique traditionally used in retail market research, to identify frequent and characteristic genotype patterns showing strong associations to phenotype clusters. We applied this strategy to three independent GWAS datasets from 2,835 phenotypically characterized patients with BD. In a discovery step, 20,882 candidate association rules were extracted. Two of these - one associated with eating disorder and the other with anxiety - remained significant in an independent dataset after robust correction for multiple testing, showing considerable effect sizes (odds ratio ~ 3.4 and 3.0, respectively). Our approach may help detect novel specific genotype-phenotype relationships in BD typically not explored by analyses like GWAS. While we adapted the data mining tool within the context of BD gene discovery, it may facilitate identifying highly specific genotype-phenotype relationships in subsets of genome-wide data sets of other complex phenotype with similar epidemiological properties and challenges to gene discovery efforts.

## Introduction

It is widely accepted that the high heritability of around 80 % for bipolar disorder (BD) is conferred by a polygenic component yet to be understood in its complexity [1,2]. Genome-wide association studies of BD have identified several genome-wide significant variants and also hinted at the existence of many more variants which fail to achieve the rigorous threshold of genome-wide significance (p<5.0e-08) but contribute to the overall variance when considered within the context of polygenicity [5,6]. However, the number of newly identified variants is far below original expectations, with limited sample sizes being one of the explanatory factors. The largest sample for a meta-analysis of GWAS of BD to date comprised nearly 64,000 participants [7]. Although this is an impressive sample size, GWAS of other phenotypes, such as adult height, have demonstrated that samples three-times this figure are required to achieve an adequate number of significant findings [8]. Recent successes of the Psychiatric Genomics Consortium (https://pgc.unc.edu/ in schizophrenia genetics where case-control samples have already exceeded 100,000 individuals suggest that continued enlargement of sample size will also increase the yield of genome-wide significant findings for BD. Clinical heterogeneity of the BD phenotype may also have hampered success in identifying vulnerability genes. DSM [9] and ICD [10] present a list of possible symptoms, each of which must persist for a minimum period of time for the diagnosis to be assigned. Since a diagnosis of BD is based upon the presence of a minimum number of these symptoms, the diagnosis can be assigned for varying symptom constellations. Thus the nature and number of the underlying clinical symptoms, as well as the time periods over which they occur, show substantial variation between patients. Thus, the clinical presentation is diverse, and differing disease courses are observed within each diagnostic category.

We hypothesize that heterogeneity can be reduced and the number of identified variants increased by analyzing the joint effect of several genetic variants on specific subsets of clinical items identified in BD patients [11,12]. We hypothesize that systematic data mining approaches from other fields can be applied to analyses of GWAS data. Popular methods such as support vector machines, Bayesian networks, and association rule mining (ARM) have been successfully applied in industry. ARM is one of the most important and well researched techniques of data mining [13]. It aims to extract casual structures among sets of items in data bases for discovering and predicting regularities and has been applied extensively to market research [14,15] in order to analyze customer habits. Recently, it has been introduced to biological data, in particular microarray data for gene expression analysis [16,17]. We consider this approach highly appropriate for genome-wide data, since its main goal is to unravel unknown associations between source data, i.e. customer profiles in market research, and potential targets, i.e. their buying behavior, which can then be used for target prediction (Figure 1). Within the context of genome-wide data, the source data are genetic variants and the potential targets are symptom clusters. The aim of the present study is to apply this data mining approach to GWAS datasets of BD in order to identify yet undetected genotype-phenotype associations, searching for associations between frequently occurring genotype combinations and symptom clusters.

**Figure 1.**
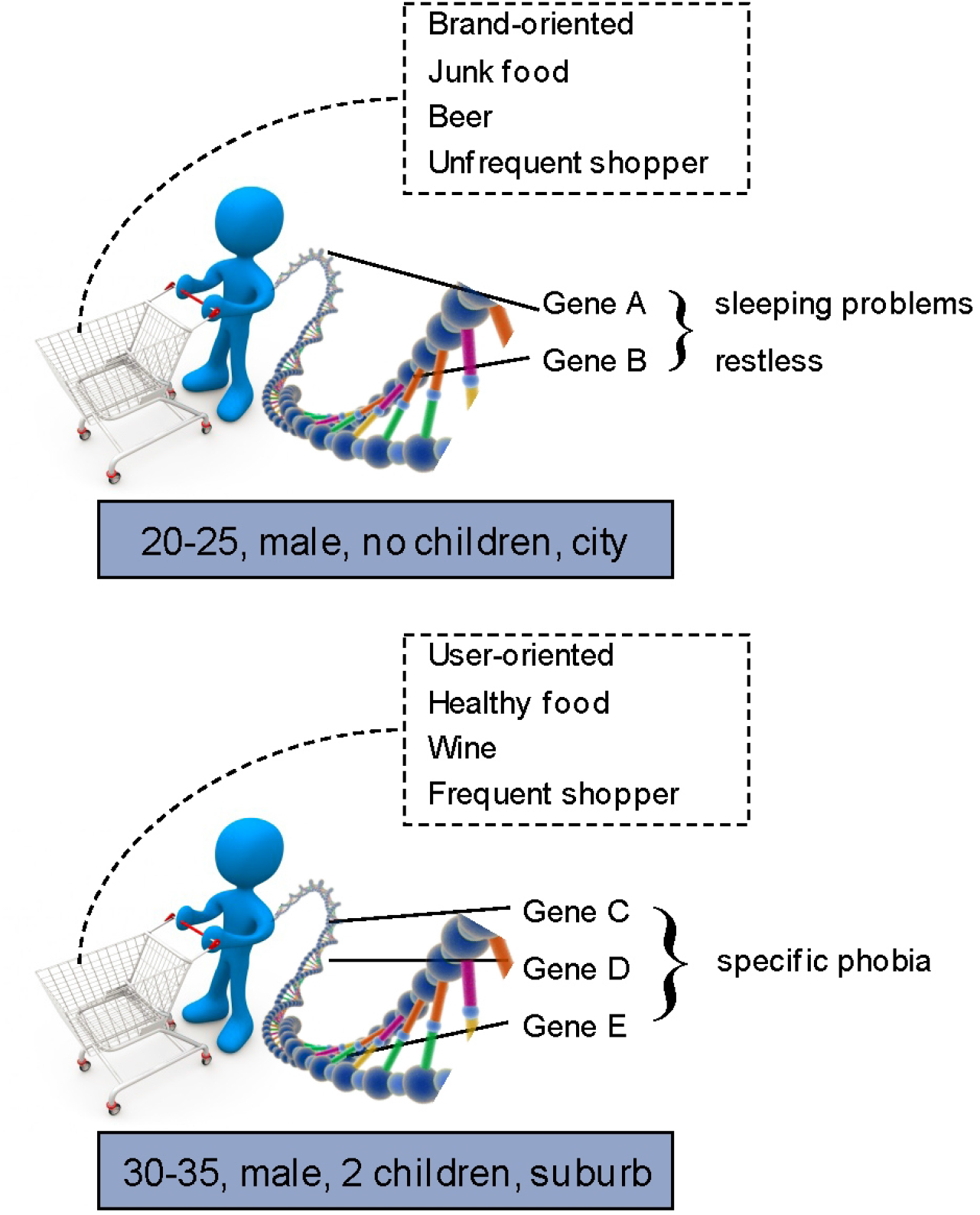
Outline of the overall approach. A main goal of market research is to identify rules that predict customer habits based on market baskets. In the cartoon, a male customer between 20 and 25 without children living in the city favours junk food and beer and when he goes shopping he will most likely buy brands. Adapting this idea to genetic research we try to identify those genetic factors from the plethora of genetic factors in the “market basket” that are characterized by specific phenotypic features (like specific phobia or restlessness). The cartoon contains graphical depictions by Benjamin Albiach Galan and Konstantinos Kokkinis.

## Materials and Methods

### Samples

Genotype and phenotype data were obtained from three independent BD case-control samples, the US-American GAIN (1,000 cases, 1,033 controls) [18] and TGEN (1,190 cases, 401 controls) collections [19] and the German BoMa (645 cases, 1310 controls) sample [20]. Clinical symptoms, sociodemographic and environmental features were ascertained using structured interviews (DIGS [21] for GAIIN and SCID-I for BoMa [22]. All phenotypes were retrieved from professionally curated databases [23,24]. Detailed information on the samples can be found elsewhere [18,19,25]. Descriptive statistics for both samples are provided in Table S1. The total sample for the present study comprised n=5,579 subjects (2,835 cases and 2,744 controls). The GAIN sample was used for the discovery step, and the TGEN and BoMa samples were used for the replication step. Prior to study inclusion, written informed consent was obtained from all subjects. The study was performed under a protocol approved by the ethical committee of the University of Heidelberg (*Medizinische Ethikkommission II*).

### Selection of clinical features

In addition to the two phenotypic specifiers age at onset (AAO) and sex, we included a variety of other phenotypic feature, for the selection of which we applied the following criteria: (i) evidence of familiality and/or heritability [26]; (ii) a frequency of at least 5% across all three samples; (iii) a missing data rate of less than 10%; (iv) availability in at least two of the three data sets; and/or (v) clinical features with a high frequency among BD patients, including comorbid features not being part of the diagnosis of BD. In total, we selected 23 clinical features (Table S2), the frequency of which was similar across all three samples (Figure S1), and ranged from <10% (e.g. eating disorder) to 80% (e.g. reckless behavior).

### Selection of single markers and genetic model

The GAIN and TGEN samples were genome-wide genotyped on the Affymetrix 6.0 SNP array. For the BoMa sample, the Illumina HumanHap550 BeadChip was used. All genotypes were imputed based on 2.1 million HapMap Phase 2 markers [27]. Due to computational runtime constraints, our analysis is based on a selected number of markers. We included only those SNPs that showed an association p-value of less than 0.001 in a recent meta-analysis of 4,961 BD patients and 7,294 controls (Supplementary Notes, Methods-SNP *selection*). Our resulting SNP set comprised 5,487 SNPs, on which LD pruning (Supplementary Notes, Methods-*Linkage disequilibrium*) was performed in order to reduce redundancy within the genotype data before the discovery step and to decrease runtime. This left us with a total of 1,599 SNPs. Of these, 1,581 SNPs were available in all three samples studied. As the ARM approach requires binary variables we had to transform the genotype information into a binary format (Supplementary Notes, Methods-*Genetic Models*).

### Algorithm for association rule mining

The basic idea for identifying genotype-phenotype data using these binary genotype data is to (i) receive frequent genotype patterns, (ii) to look for significantly associated phenotypes as candidates, or in terms of the original algorithm *candidate association rules*, in a discovery dataset, and (iii) to validate these candidate association rules in an independent replication dataset. Figure 2 illustrates the basic idea of combining genotypic information in order to identify frequent genotype patterns (left) and evaluate the patterns regarding interesting phenotype traits in order to receive a candidate association rule like genotype-pattern A implies phenotype-pattern B (*A* ⇒ *B*) (right).

**Figure 2.**
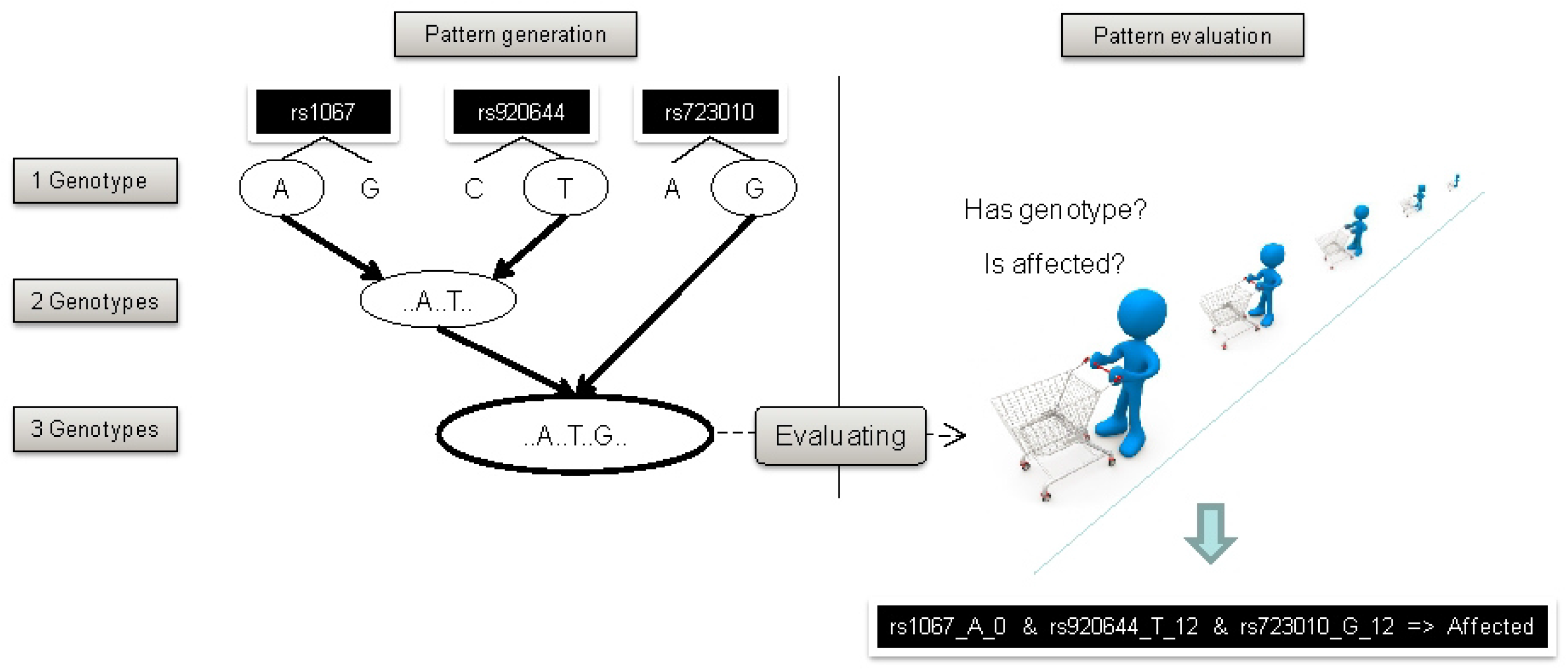
Illustration of the implemented version of the association rule mining algorithm. The lattice shown left is traversed starting from root { } to all leaves. Each genotype pattern (node in the tree) represents a subgroup of patients shown in the genotype matrix G. Additionally, using the p phenotype information of the patients from matrix P, we can count genotype and phenotype occurrences in contingency tables. Here illustrated for the genotype pattern g_1_g_2_g_n_ with ‘a’ counting all patients where genotype g_1_g_2_g_n_ and phenotype p_i_ are present, ‘d’ were neither of both are present, and ‘b’ and ‘c’ counting patients with presentation of genotype g1g2gn but not phenotype pi and visa versa. The lattice is traversed as long as there are unprocessed genotype patterns that cannot be pruned before.

#### Identifying frequent genotype patterns

The frequent genotype patterns can be identified in a systematic manner. Several approaches have been developed for association rule mining [28,29]. Here we use the most common Apriori algorithm, as it can be implemented in a straightforward manner and shows a good performance for short patterns, making it an ideal choice for the present study. For details, see Supplementary Notes, Methods-*Runtime*, Methods-*Apriori algorithm*, and Methods-*Closed frequent itemsets*.

#### Discovery of candidate association rules

Once a frequent genotype pattern is identified it is tested for association with each phenotypic trait, i.e. each of the 23 selected clinical features. This step involves the generation of a contingency table for the frequent genotype pattern and each clinical feature. Based on this contingency table, the interestingness of an association rule is assessed. For details, see Supplementary Notes, *Methods-Association rule discovery.*

#### Replication of candidate association rules

The third stage of our rule mining approach is the replication of the candidate rules. Significance testing is rarely investigated in rule mining [30]. However, we considered this to be important as an inherent aspect of rule mining is the occurrence of false positive results. We anticipated 50,000 false positives per one million tests on the basis of the widely used type I error rate of 5%. One approach to correct for this is to assume that the association rules are independent and apply Bonferroni correction to the test statistics in the discovery data only. However, several simulations have shown that as well as reducing the rate of false discoveries, the Bonferroni approach also reduces the rate of true positive findings [30]. Alternatively, permutation tests can be performed to test whether or not the association between a genotype pattern and a phenotype cluster is random (Supplementary Notes, Methods-*Permutation tests*). However, when constrained to a single dataset, both methods are susceptible to overfitting. Thus, we considered the performance of a replication of all candidate rules nCR in an independent dataset a more appropriate alternative as this adjusts for potentially spurious sample effects and random associations. Using the latter approach and the Bonferroni method, we defined a primary test-wide significance level *α_adj_* for the replication as:

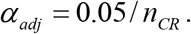

However, as shown by our findings and those of Webb [30], when extracting a set of association rules using the ARM approach, the rules are unlikely to be independent. Thus, this significance testing remains conservative and is likely to reject true positive results. Therefore, we also report p-values adjusted using the false discovery rate (FDR). This is an alternative statistical method to adjust for multiple testing: FDR assumes sub-groups of tests to be dependent. FDR is less conservative, resulting in an increase in power at the cost of an increased likelihood of type I errors [31,32].

#### Analyses

In order to apply our approach to the GWAS data, we developed a software tool, termed RUDI (**RU**le **Di**scoverer; Supplementary Notes, Software). A rule discovery analysis of 1,581 SNPs (3,162 variables) and 23 phenotypic traits in the 1,000 cases from the GAIN sample was performed. Around 4.286e+09 genotype patterns were tested using the following settings: (a) z-score of 5.0; (b) maximum length of the genotype pattern of 3, and (c) absolute minimum support of individuals matching the particular genotype pattern of 50 (Supplementary Notes, Methods-Parameter *selection*). The runtime on an Intel Xeon X3220 with 2.4GHz was around 18 hours on a single processor using the described settings. A second run was performed to replicate the candidate rules in our replication dataset of n=1,835 BD patients (TGEN + BoMa).

## Results

N=20,882 candidate rules satisfied the required thresholds in the discovery data set. The strongest association rule showed a p-value of 3.457e-15 (#962) and thus reached significance after correction for multiple testing using the Bonferroni method (adjusted p-value=3.260e-04). When all candidate rules were compared, 15 disjunct phenotype clusters were observed. Of these, 11 consisted of a single clinical feature. The remaining four consisted of two clinical features (Table S3).

Replication of the n=20,882 candidate rules was then performed in our replication dataset of n=1,835 BD patients. The level of significance after adjustment using the Bonferroni method was 2.394e-06 for a default alpha of 5%. Although replication of the top finding from the discovery step (#962) failed, two rules met the significance threshold:

i. rule #12978

> rs6733011_A_0, rs4113925_T_0, rs3769745_T_0 => ‘eating disorder’ (ED)

with a p-value = 3.576e-08 and an odds ratio (OR) = 3.566 [.95 confidence interval (CI): 2.169-5.681];
and
ii. rule #6221

> rs858057_G_0, rs4757144_G_0, rs3130781_C_0 => ‘simple phobia’ (SP)

with a p-value = 1.780e-06 and an OR = 2.995 [.95 CI: 1.841-4.730]. Three further rules remained significant after FDR correction (Supplementary Notes, *Further results*). A total of 1,252 (6.0%) of the candidate rules reached nominal significance in the replication sample. The distribution of the p-values for all candidate rules in the replication dataset fits the expected chi-squared distribution (Figure S2).

### Association finding with eating disorder

Our top finding, rule #12978, showed a genotype pattern frequency of 5.2-7.4% in the case samples and of around 5.2-5.8% in the control populations (Table S4). Further details of the genotype pattern are shown in Table S7. In addition to the primary replication within the discovery-replication framework, two types of permutation tests (Supplementary Notes, *Methods-Permutation tests*) were performed to estimate: (a) the probability of finding a more significant association with the genotype pattern by re-sampling the phenotype; and (b) the probability of randomly choosing a genotype pattern that shows at least the same level of significance. Both reject the hypothesis of a random association based on the empirical p-values observed (4.000e-06 and 7.000e-06, respectively), based on 1e+06 trials in the discovery data. In a subsequent step, we combined the data of all n=2,835 patients and reevaluated rule #12978, i.e. we compared patients with and without an eating disorder (BD_ED and BD_nonED, respectively) in terms of the genotype pattern of this rule. This combined analysis of cases showed a p-value=5.300e-14 and an OR=4.120 [.95 CI: 2.740-6.068]. Thus, within the group of patients carrying the genotype pattern, the frequency of a co-morbid eating disorder is increased on average by a factor of 4. An association analysis was then performed for each of the three SNPs of the genotype pattern to determine whether the observed association was due to the combination of the three SNPs or conferred by only one of them. Single trend tests for the phenotype ‘eating disorder’ was performed in each of the three datasets using PLINK [33]. No significant evidence was found to support the hypothesis that the association with the phenotype of the rule is driven by a single SNP (Table S6).

We furthermore performed an association study of the genotype pattern of rule #12978 in cases versus controls. No differential distribution of the genotype pattern was observed between (a) the BD_nonED cases and controls and (b) between all BD cases and controls: However, the genotype pattern was significantly associated with BD_ED cases compared to controls (p-value=4.937e-14, OR=4.107 [.95 CI: 2.735-6.040]) (Table S5).

### Association finding with simple phobia

The second finding, rule #6221, showed an association with ‘simple phobia’. The genotype frequencies were 5.4-6.9% in cases and 5.1-7.2% in controls. In the combined analysis of all cases, we observed a p-value=3.476e-13 (adjusted p-value=3.427e-02) and an OR=3.551 [.95 CI: 2.453-5.063]. Thus, within the group of patients carrying the genotype pattern, the frequency of a co-morbid simple phobia increased on average by a factor of 3.5. As was the case for the rule including eating disorder, the association was not conferred by the single SNPs taken separately but only in combination (Table S6). Likewise a case-control analysis showed: (a) no differential distribution of the genotype pattern between the BD_nonSP cases and controls nor (b) between all BD cases and controls. However, we observed (c) a significant differential distribution between BD_SP cases and controls (p-value=1.686e-11, OR=3.195 [.95 CI: 2.220-4.523]) (Table S5).

## Discussion

Application of the ARM data mining approach identified significant associations between sets of candidate SNPs and BD subgroups characterized by two specific comorbid conditions: eating disorder and simple phobia.

Our top finding (rule #12978) highlights an association between the genotype pattern of rule #12978 and the subgroup of BD patients with an eating disorder. The association was conferred by the combination of three SNPs but not by the individual SNPs. While the proportion of BD patients with an eating disorder was very small (n=192 patients; i.e. 6.8% of our sample), this frequency is comparable to that reported in other studies [34, 35]. Thirty-seven of these patients displayed the genotype pattern of rule #12978, which was present in 182 of all BD patients. Despite the small sample size, the association finding (p-value=4.937e-14) is rather strong with an OR=4.107 in the combined case-control analysis, an effect size typically not seen for diagnosis-based studies.

The likelihood that our findings may be due to chance is further decreased when considering the following two points: Firstly, the replication sample was comprised of two smaller samples, and in both of these samples, the effect was in the same direction (with test-wide significance being achieved in neither). Secondly, our findings fall in line with reports on the function of the genes involved. SNP rs3769745 of rule #12978 is located in the intron region of the cyclic nucleotide gated channel alpha 3 gene (*CNGA3*) on chromosome 2. In humans, *CNGA3* is implicated in total color blindness (achromatopsia) [36,37]. Animal studies have shown that *CNGA3* is required for normal vision [38], olfactory signal transduction [39], and involved in nociceptive processing [40]. Further, it is expressed in the mouse brain and is reported to influence synaptic plasticity and behaviour [41]. Research has also shown that the specialized olfactory subsystem to which CNGA3 belongs is required for the acquisition of socially transmitted food preferences (STFPs) in mice. Mice that lack this gene fail to acquire STFPs from other mice, and exhibit an absence of neuronal activation of the ventral subiculum of the hippocampus, a brain region implicated in STFP retrieval [42]. According to the KEGG Database, *CNGA3* is in a common pathway, .e., olfactory transduction (KEGG ID hsa04740), with *CALM1*, a candidate gene for anorexia nervosa [43]. To the best of our knowledge, no association between this variant and eating disorder has been reported so far. For the other two variants, a plausible support from biological data is not available. SNP rs6733011 is located in an intron region of the KIAA1211-like (*KIAA1211L*) gene on chromosome 2 that encodes the uncharacterized protein *C2orf55* (chromosome 2 open reading frame 55). The location is within a 500 kb window to rs3769745, but not in the same LD block (r^2^ = 0.027 and D’ = 0.343 in the discovery dataset). Its function remains unknown. SNP rs4113925 is located on chromosome 12q24.21 in an intron of the T-box transcription factor (*TBX5*) gene. This T-box gene has been implicated in heart development and disease as well as specification of limb identity [44].

To investigate whether our finding identified genetic markers specific to BD with an eating disorder subphenotype or eating disorder per se, we tested a potential association of the genotype pattern of rule #12978 with an eating disorder phenotype comprising anorexia and bulimia in a population-based sample from Australia (n=1672, 12.9 % with a diagnosis of anorexia or bulimia). We did not see an association of the genotype pattern of rule #12978, suggesting that our approach has detected a genetic marker for BD with comorbid eating disorder rather than for eating disorder per se.

Our second finding, rule #6221, showed an association with simple phobia. Two of the three contributing SNPs are located within genes. SNP rs4757144 is located in an intron region of the aryl hydrocarbon receptor nuclear translocator-like (*ARNTL*) gene, and rs3130781 is located in an intron region of the diffuse panbronchiolitis critical region 1 (*DPCR1*) gene. The third SNP, rs858057, is located at an intergenic region of 20p11.21. An implication of ARNTL in the etiology disorders via its influence on the circadian system has been discussed repeatedly. 45–50], There is further report that genes homologous to *ARNTL* may be implicated in the etiology of anxiety. Sipilä and colleagues [50] tested several anxiety phenotypes for association with 13 circadian genes and found association between social phobia and *ARNTL2.* Thus the *ARNTL* gene family may be involved in this co-morbid phenotype. The second gene, *DPCR1*, is located in the major histocompatibility complex (MHC), which hosts genes that are crucial for the functioning of the immune system.

While we observed several other genotype-phenotype rules that may warrant further in-depth investigation (Table 1 and Supplementary Notes, *Further results*), we focused on the rules implicating BD subtypes with comorbid eating disorder and simple phobia, respectively, as only these two survived our stringent multi-tiered evaluation of potential type I error. These steps help minimize-if not eliminate-type I error rate in ARM due to the overfitting of rules in a particular dataset [28].

**Table 1.**
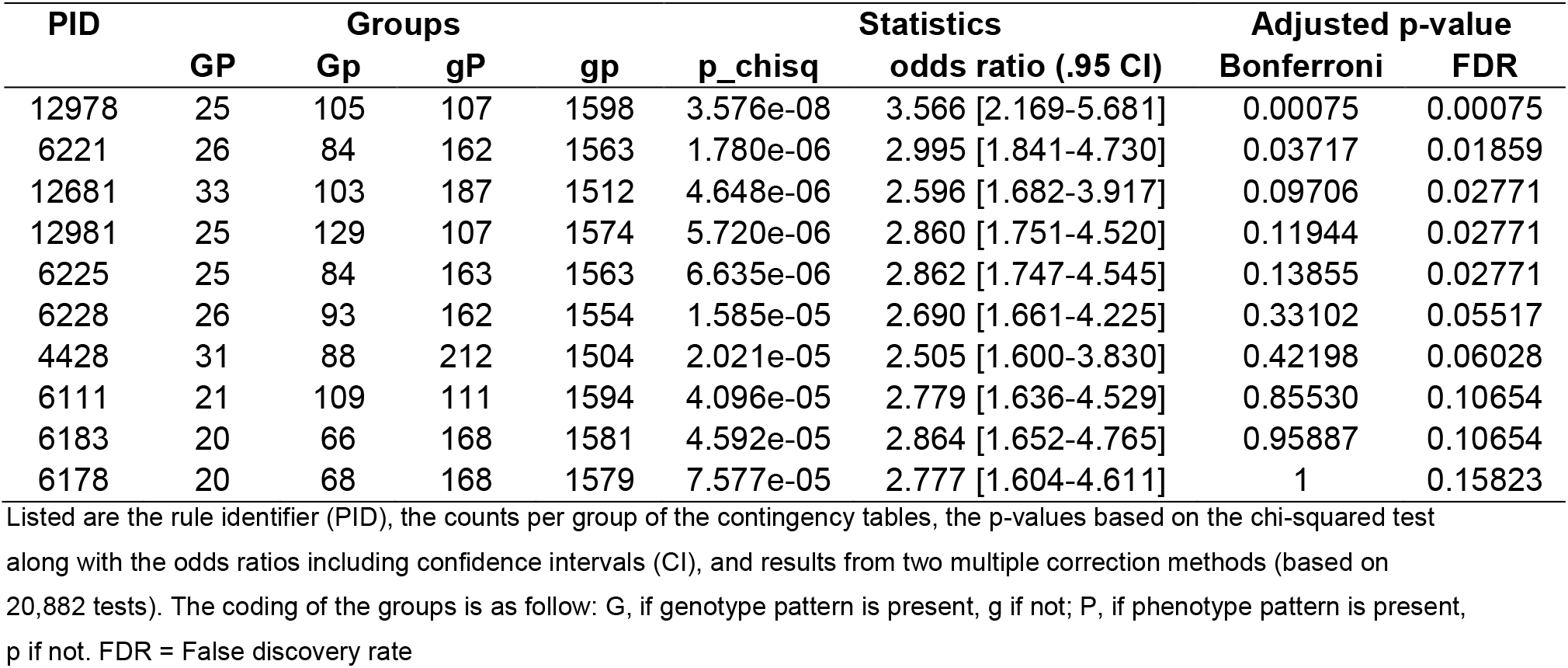
Top 10 association rules regarding their p-values in the replication dataset (TGEN+BoMa).

We would like to point out that the two reported association rules were associated with very low frequency phenotypes. This is due to the characteristics of the z-score approach applied. Since small proportions of the data are more likely to deviate strongly from the random distribution, larger effects and thus larger z-scores are expected. As only those rules that show a z-score of greater or equal 5 are extracted as candidates, this particular rule measure is biased towards associations with low frequency phenotypes. This further explains the relatively small number, i.e. 4 out of 15 (Supplementary Notes, Methods-*Phenotype cluster*), of phenotype clusters that consisted of more than one phenotype. We may thus have discarded many potentially true findings. Given that our study can be considered a proof-of-concept for the application of ARM on GWAS-derived data for a complex phenotype, we opted for statistical stringency rather than a merely exploratory pattern mining. While this approach resulted in only two findings, they are characterized by effect sizes up to four times larger than typically seen in GWAS of BD or other complex traits.

In summary, using already available GWAS data sets on BD, we have established and implemented a novel data mining process for complex genetic data. We identified genotype-phenotype patterns highlighting subtypes of BD characterized by specific comorbid conditions. These two comorbid conditions, eating disorder and simple phobia, may delineate more homogenous subgroups of BD that warrant further study in genomic studies of BD. An important limitation of our approach is that our approach was only based on 5487 SNPs that showed some evidence of association with BD. As association rule mining may detect hidden association of specific phenotypes with previously un-identified SNPs, our approach may have missed several novel associations. This restriction, however, was due to our motivation to perform genotype-phenotype dissection on SNPs that showed some evidence of association. We were further bound by some computational runtime constraints. Further extensions of the algorithm will be required to allow for a variety of assumed genetic models (here we used a dominant genetic model), to optimize computational feasibility for an increased number of SNPs (Supplementary Notes, Methods-Runtime), and to determine the optimal correction methodology for highly correlated data

Our approach highlights a strategy for genotype-phenotype dissection and for the identification of genetic susceptibility variants beyond initial GWAS of heterogeneous disorders. Finally, our results emphasize the importance of thorough phenotyping, particularly with regard to comorbidity.

## Acknowledgments

We gratefully acknowledge critical input from Nicholas Martin, Scott Gordon, and Cynthia. This study is part of the *Systematic Investigation of the Molecular Causes of Major Mood Disorders and Schizophrenia* (*MooDS*) network, which is funded by the Federal Ministry of Education and Research (BMBF) through the framework of National Genomic Research Network (NGFN) (Grant 01GS08144 to SC and MMN; Grant 01GS08147 to MR). MR was also supported by the Seventh Framework Program of the European Union (FP7/2007-2011) under grant agreement no. 242257 (ADAMS). MMN also received support from the Alfried Krupp von Bohlen und Halbach-Stiftung. LK, FJM, and TGS were also supported through Intramural Research Program of the National Institute of Mental Health (NIHM) at the National Institutes of Health (NIH) of the US Government.

## References

1. McGuffin P, Rijsdijk F, Andrew M, Sham P, Katz R, et al. (2003) The heritability of bipolar affective disorder and the genetic relationship to unipolar depression. Arch Gen Psychiatry 60: 497–502. doi:10.1001/archpsyc.60.5.497.

2. Craddock N, O’Donovan MC, Owen MJ (2005) The genetics of schizophrenia and bipolar disorder: dissecting psychosis. J Med Genet 42: 193–204. doi: 10.1136/jmg.2005.030718.

3. Lee KW, Woon PS, Teo YY, Sim K (2012) Genome wide association studies (GWAS) and copy number variation (CNV) studies of the major psychoses: what have we learnt? Neurosci Biobehav Rev 36: 556–571. doi:10.1016/j.neubiorev.2011.09.001.

4. Sullivan PF, Daly MJ, O’Donovan M (2012) Genetic architectures of psychiatric disorders: the emerging picture and its implications. Nature Reviews Genetics 13: 537–551. doi: 10.1038/nrg3240.

5. Lee SH, DeCandia TR, Ripke S, Yang J; Schizophrenia Psychiatric Genome-Wide Association Study Consortium (PGC-SCZ); International Schizophrenia Consortium (ISC); Molecular Genetics of Schizophrenia Collaboration (MGS), Sullivan PF, Goddard ME, Keller MC, Visscher PM, Wray NR (2012) Estimating the proportion of variation in susceptibility to schizophrenia captured by common SNPs. Nat Genet 44:247–250

6. Schulze TG, Akula N, Breuer R, Steele J, Nalls MA, Singleton AB, Degenhardt FA, Nöthen MM, Cichon S, Rietschel M; Bipolar Genome Study, McMahon FJ (2014) Molecular genetic overlap in bipolar disorder, schizophrenia, and major depressive disorder. World J Biol Psychiatry 15:200–208.

7. Sklar P, Ripke S, Scott LJ, Andreassen OA, Cichon S, et al. (2011) Large-scale genome-wide association analysis of bipolar disorder identifies a new susceptibility locus near ODZ4. Nat Genet 43: 977–983. doi:10.1038/ng.943.

8. Lango Allen H, Estrada K, Lettre G, Berndt SI, Weedon MN, et al. (2010) Hundreds of variants clustered in genomic loci and biological pathways affect human height. Nature 467: 832–838. doi:10.1038/nature09410.

9. American Psychiatric Association (2000) Diagnostic and statistical manual of mental disorders: DSM-IV-TR. Washington, DC: American Psychiatric Association.

10. World Health Organization (2011) International statistical classification of diseases and related health problems. Geneva: World Health Organization.

11. Purcell SM, Wray NR, Stone JL, Visscher PM, O’Donovan MC, et al. (2009) Common polygenic variation contributes to risk of schizophrenia and bipolar disorder. Nature 460: 748–752. doi:10.1038/nature08185.

12. Lee SH, Wray NR, Goddard ME, Visscher PM (2011) Estimating missing heritability for disease from genome-wide association studies. Am J Hum Genet 88: 294–305. doi:10.1016/j.ajhg.2011.02.002.

13. Kotsiantis S, Kanellopoulos D (2006) Association Rules Mining: A Recent Overview. International Transactions on Computer Science and Engineering: pp. 71–82.

14. Agrawal R, Imielinski T, Swami A (1993) Mining Association Rules between Sets of Items in Large Databases. Proceedings of the 1993 ACM SIGMOD International Conference on Management of Data. 2. Washington DC, USA: ACM Press, Vol. 22. pp. 207–216.

15. Ngai EWT, Xiu L, Chau DCK (2009) Application of data mining techniques in customer relationship management: A literature review and classification. Expert Systems with Applications 36: 2592–2602. doi:10.1016/j.eswa.2008.02.021.

16. Martinez R, Pasquier N, Pasquier C (2008) GenMiner: mining non-redundant association rules from integrated gene expression data and annotations. Bioinformatics 24: 2643–2644. doi:10.1093/bioinformatics/btn490.

17. Liu Y-C Cheng C-P, Tseng VS (2011) Discovering relational-based association rules with multiple minimum supports on microarray datasets. Bioinformatics 27: 3142–3148. doi:10.1093/bioinformatics/btr526.

18. Smith EN, Bloss CS, Badner JA, Barrett T, Belmonte PL, et al. (2009) Genome-wide association study of bipolar disorder in European American and African American individuals. Mol Psychiatry 14: 755–763. doi:10.1038/mp.2009.43.

19. Smith EN, Koller DL, Panganiban C, Szelinger S, Zhang P, et al. (2011) Genome-wide association of bipolar disorder suggests an enrichment of replicable associations in regions near genes. PLoS Genet 7: e1002134. doi:10.1371/journal.pgen.1002134.

20. Cichon S, Mühleisen TW, Degenhardt FA, Mattheisen M, Miró X, et al. (2011) Genome-wide association study identifies genetic variation in neurocan as a susceptibility factor for bipolar disorder. Am J Hum Genet 88: 372–381. doi:10.1016/j.ajhg.2011.01.017.

21. Nurnberger JI Jr, Blehar MC, Kaufmann CA, York-Cooler C, Simpson SG, et al. (1994) Diagnostic interview for genetic studies. Rationale, unique features, and training. NIMH Genetics Initiative. Arch Gen Psychiatry 51: 849–859; discussion 863-864.

22. Spitzer RL, Williams JB, Gibbon M, First MB (1992) The Structured Clinical Interview for DSM-III-R (SCID). I: History, rationale, and description. Arch Gen Psychiatry 49: 624–629.

23. Potash JB, Toolan J, Steele J, Miller EB, Pearl J, et al. (2007) The bipolar disorder phenome database: a resource for genetic studies. Am J Psychiatry 164: 1229–1237. doi:10.1176/appi.ajp.2007.06122045.

24. Fangerau H, Ohlraun S, Granath RO, Nöthen MM, Rietschel M, et al. (2004) Computer-Assisted Phenotype Characterization for Genetic Research in Psychiatry. Human Heredity 58: 122–130. doi:10.1159/000083538.

25. Baum AE, Akula N, Cabanero M, Cardona I, Corona W, et al. (2008) A genome-wide association study implicates diacylglycerol kinase eta (DGKH) and several other genes in the etiology of bipolar disorder. Mol Psychiatry 13: 197–207. doi: 10.1038/sj.mp.4002012.

26. Schulze TG (2006) What is familial about familial bipolar disorder? Resemblance among relatives across a broad spectrum of phenotypic characteristics. Archives of General Psychiatry 63: 1368. doi:10.1001/archpsyc.63.12.1368.

27. McMahon FJ, Akula N, Schulze TG, Muglia P, Tozzi F, et al. (2010) Meta-analysis of genome-wide association data identifies a risk locus for major mood disorders on 3p21.1. Nat Genet 42: 128–131. doi:10.1038/ng.523.

28. Han J, Kamber M (2006) Data mining concepts and techniques, second edition. 2nd ed. Amsterdam; Boston; San Francisco Calif.: Elsevier; Morgan Kaufmann Publishers.

29. Maimon OZ, Rokach L (2005) Data mining and knowledge discovery handbook. New York, NY, USA: Springer.

30. Webb GI (2006) Discovering significant rules. Proceedings of the 12th ACM SIGKDD international conference on Knowledge discovery and data mining. New York, NY, USA: ACM Press. pp. 434–443. doi:10.1145/1150402.1150451.

31. Benjamini Y, Hochberg Y (1995) Controlling the False Discovery Rate: A Practical and Powerful Approach to Multiple Testing. Journal of the Royal Statistical Society, Series B (Methodological): 289–300.

32. Benjamini Y (2010) Simultaneous and selective inference: Current successes and future challenges. Biom J 52: 708–721. doi:10.1002/bimj.200900299.

33. Purcell S, Neale B, Todd-Brown K, Thomas L, Ferreira MAR, et al. (2007) PLINK: a tool set for whole-genome association and population-based linkage analyses. Am J Hum Genet 81: 559–575. doi:10.1086/519795.

34. McElroy SL, Kotwal R, Keck PE Jr (2006) Comorbidity of eating disorders with bipolar disorder and treatment implications. Bipolar Disord 8: 686–695. doi: 10.1111/j.1399- 5618.2006.00401.x.

35. McElroy SL, Frye MA, Hellemann G, Altshuler L, Leverich GS, et al. (2011) Prevalence and correlates of eating disorders in 875 patients with bipolar disorder. Journal of Affective Disorders 128: 191–198. doi:10.1016/j.jad.2010.06.037.

36. Ding X-Q, Fitzgerald JB, Quiambao AB, Harry CS, Malykhina AP (2010) Molecular pathogenesis of achromatopsia associated with mutations in the cone cyclic nucleotide-gated channel CNGA3 subunit. Adv Exp Med Biol 664: 245–253. doi:10.1007/978-1- 4419-1399-9_28.

37. Lam K, Guo H, Wilson GA, Kohl S, Wong F (2011) Identification of variants in CNGA3 as cause for achromatopsia by exome sequencing of a single patient. Arch Ophthalmol 129: 1212–1217. doi:10.1001/archophthalmol.2011.254.

38. Biel M, Seeliger M, Pfeifer A, Kohler K, Gerstner A, Ludwig A, Jaissle G, Fauser S, Zrenner E, Hofmann F (1999) Selective loss of cone function in mice lacking the cyclic nucleotide-gated channel CNG3. Proc Natl Acad Sci U S A 96:7553–7557.

39. Leinders-Zufall T, Cockerham RE, Michalakis S, Biel M, Garbers DL, Reed RR, Zufall F, Munger SD (2007) Contribution of the receptor guanylyl cyclase GC-D to chemosensory function in the olfactory epithelium. Proc Natl Acad Sci U S A 4; 104(36):14507–14512.

40. Heine S, Michalakis S, Kallenborn-Gerhardt W, Lu R, Lim HY, Weiland J, Del Turco D, Deller T, Tegeder I, Biel M, Geisslinger G, Schmidtko A (2011) CNGA3: a target of spinal nitric oxide/cGMP signaling and modulator of inflammatory pain hypersensitivity. J Neurosci 31:11184–11192.

41. Michalakis S, Kleppisch T, Polta SA, Wotjak CT, Koch S, et al. (2011) Altered synaptic plasticity and behavioral abnormalities in CNGA3-deficient mice. Genes Brain Behav 10: 137–148. doi:10.1111/j.1601-183X.2010.00646.x.

42. Munger SD, Leinders-Zufall T, McDougall LM, Cockerham RE, Schmid A, et al. (2010) An olfactory subsystem that detects carbon disulfide and mediates food-related social learning. Curr Biol 20: 1438–1444. doi:10.1016/j.cub.2010.06.021.

43. Pinheiro AP, Bulik CM, Thornton LM, Sullivan PF, Root TL, et al. (2010) Association study of 182 candidate genes in anorexia nervosa. Am J Med Genet B Neuropsychiatr Genet 153B: 1070–1080. doi:10.1002/ajmg.b.31082.

44. Wang C, Cao D, Wang Q, Wang D-Z (2011) Synergistic activation of cardiac genes by myocardin and Tbx5. PLoS ONE 6: e24242. doi:10.1371/journal.pone.0024242.

45. Mansour HA, Wood J, Logue T, Chowdari KV, Dayal M, et al. (2006) Association study of eight circadian genes with bipolar I disorder, schizoaffective disorder and schizophrenia. Genes Brain Behav 5: 150–157. doi:10.1111/j.1601-183X.2005.00147.x.

46. Le-Niculescu H, Patel SD, Bhat M, Kuczenski R, Faraone SV, et al. (2009) Convergent functional genomics of genome-wide association data for bipolar disorder: Comprehensive identification of candidate genes, pathways and mechanisms. American Journal of Medical Genetics Part B: Neuropsychiatric Genetics 150B: 155–181. doi:10.1002/ajmg.b.30887.

47. Nakatani N (2006) Genome-wide expression analysis detects eight genes with robust alterations specific to bipolar I disorder: relevance to neuronal network perturbation. Human Molecular Genetics 15: 1949–1962. doi:10.1093/hmg/ddl118.

48. Nievergelt CM, Kripke DF, Barrett TB, Burg E, Remick RA, et al. (2006) Suggestive evidence for association of the circadian genes PERIOD3 and ARNTL with bipolar disorder. Am J Med Genet B Neuropsychiatr Genet 141B: 234–241. doi:10.1002/ajmg.b.30252.

49. Shi J, Wittke-Thompson JK, Badner JA, Hattori E, Potash JB, et al. (2008) Clock genes may influence bipolar disorder susceptibility and dysfunctional circadian rhythm. American Journal of Medical Genetics Part B: Neuropsychiatric Genetics 147B: 1047–1055. doi:10.1002/ajmg.b.30714.

50. Sipilä T, Kananen L, Greco D, Donner J, Silander K, et al. (2010) An association analysis of circadian genes in anxiety disorders. Biol Psychiatry 67: 1163–1170. doi:10.1016/j.biopsych.2009.12.011.

